# *Candida albicans* infection suppresses Lipopolysaccharide or *Pseudomonas aeruginosa* stimulated murine bone marrow derived macrophage (BMDM) responses

**DOI:** 10.1101/2025.06.05.657992

**Authors:** Christa P. Baker, Stephanie Laba, Jordan Warner, Karen Shepherd, Heather M. Wilson, J. Simon C. Arthur

## Abstract

*Candida albicans* is a commensal fungus which populates most healthy individuals’ microbiota but can turn opportunistic in immunocompromised individuals and cause severe disease linked with high rates of mortality. With limited therapeutic options and increasing resistance to antifungals, novel treatment strategies for *C. albicans* infections is paramount. The exact immune response to *C. albicans* infections can be influenced by the surrounding microenvironment, for example, metabolic stresses or co-infection; although, knowledge on whether responses are enhanced or inhibited is lacking. Macrophages are a key immune cell in defence against *C. albicans* infection through phagocytic uptake and cytokine production that alerts other immune defence mechanisms. Here, we utilise a discovery screen approach using Data Independent Acquisition (DIA) based total proteomics to describe murine bone marrow derived macrophage (BMDM) response to *C. albicans* infection as well as in response to co-infection with gram-negative bacterial outer membrane component lipopolysaccharide (LPS) or live gram-negative bacteria *Pseudomonas aeruginosa*. We found *C. albicans* induced a surprisingly muted immune response in BMDMs as compared to LPS or *P. aeruginosa*. Moreover, upon co-infection with LPS or *P. aeruginosa, C. albicans* suppressed the BMDM proteome landscape and selectively suppressed BMDM secreted IL-6 and IL-12p40 cytokine responses to *P. aeruginosa*. Thus, *C. albicans* has significant suppressive capabilities in the host innate immune responses that could impact clinical outcomes during infection.

**Author Summary:** Human fungal pathogens are of increasing concern for global health due to infections in hospitalised patients, limited treatment options and increased resistance to antifungal treatments. The host inflammatory response to invasive infections like *Candida albicans* are important for trying to contain the pathogen. Pathogen evasion of this immune response poses a critical threat and can increase host morbidity and mortality. Macrophages are an important innate immune cell which recognises and responds to *C. albicans* infection *in vivo*. Here, we found that *in vitro* macrophage signalling, secreted cytokines and the total proteomic responses to *C. albicans* infection was much more subtle than the macrophage responses to live bacterial infection or the bacterial outer membrane component, lipopolysaccharide (LPS). Co-infection of *C. albicans* and bacteria suppressed selective macrophage protein expression including important inflammatory cytokines, IL-6 and IL-12p40. Thus, we describe an impressive suppressive response by *C. albicans* which poses a potential mechanism of enhanced immune evasion.

## Introduction

Human fungal infections are increasing in incidence globally, contributing to 3.8 million deaths annually (1). One of the main contributors to fatal fungal infections are *Candida* species, including *Candida albicans.* As an opportunistic, polymorphic pathogen*, C. albicans* frequently causes mucosal infections in its yeast form which are normally cleared by the immune system in healthy individuals. In some cases, *C. albicans* may cause systemic infection. This occurs most commonly in immunocompromised individuals, and are much more serious and carry a high mortality rate (2). Globally, 1,565,000 individuals are reported to develop invasive Candidiasis, of which 63.6% individuals succumb to the infection (1). Infection with *Candida* species is also problematic in clinical settings, with Candidemia being the fourth most common hospital acquired bloodstream infection (3–6). A growing concern in the context of *Candida* infections is the increased resistance of some *Candida* species to existing antifungals drugs. This is compounded by the low number of antifungal drugs available, meaning that effective treatments are becoming increasingly problematic (7–11). Thus, new treatment strategies are critical to address the issue of rising incidence in fungal infections and their resistance to current anti-fungal drugs.

Systemic *Candida* infections can occur when the body’s cutaneous or mucosal barriers fail, allowing dissemination of *Candida* through the bloodstream as a yeast. Once in the bloodstream, *C. albicans* yeast can morphologically change into filamentous hyphae due to environmental cues including factors in the serum and temperatures of 37 °C (12). This can allow *C. albicans* hyphae to grow out from blood vessels into surrounding tissues, resulting in tissue damage and potentially organ failure and mortality (2,13,14). This complex pathogenesis mechanism requires an equally complex immune response to counter it with important roles for both innate and adaptive immunity (15–17). Macrophages play an important role in the initial response to *C. albicans* infection, as demonstrated by an increased susceptibility in *Candida* infection models following experimental depletion of various macrophage populations *in vivo* (18–20). Pattern recognition Receptors (PRRs) expressed on macrophages are able to identify pathogen associated molecular patterns (PAMPs), including structural components of *C. albicans* cell wall (21–31). Activation of these receptors can potentially stimulate the macrophage to produce inflammatory mediators to aid in pathogen killing or coordinate downstream immune reactions. PRRs together with phagocytic receptors also coordinate the engulfment of *C. albicans* through phagocytosis to aid in killing of the *Candida* (32,33). Commonly studied PRRs in the context of *Candida* infection include C-type lectin receptors (CTLRs) Dectin-1, Dectin-2, Clec4d and Clec4e, which identify both yeast and hyphal cell wall structural components β-glucan, α-mannan and mannose, respectively (21,24,25,27,34–36). Additionally, Toll-like receptors (TLRs) also identify and respond to *C. albicans* cell wall components and genetic content, also inducing an inflammatory response, including TLR1, TLR2, TLR4 and TLR6 (23,29,30,37).

Much of what we understand about the macrophage signalling response to *C. albicans* is through studies utilising isolated PRR agonists including fungal cell wall components like zymosan—from *Saccharomyces cerevisiae*—or heat killed *C. albicans* in cell culture experiments (32,38). Activation of TLRs and CTLRs results in activation of pathways which are dependent on MyD88/TRIF and Syk-CARD9 signalling cascades, respectively. Despite having different upstream components, these pathways converge on activation of ERK1/2, p38 and canonical NFκB signalling pathways, which together result in induction of pro-inflammatory cytokines including IL-6, IL-12 and TNF as well as the anti-inflammatory cytokine IL-10 (32,39–42). While these agonists are critical for understanding specific PRR responses, they may not best replicate responses to live pathogens as some pathogens, like *C. albicans, can* have outer-cell wall components which mask identification by macrophages (43,44). Studying the host response to live pathogen is intrinsically more complicated by *C. albicans* polymorphic phenotypes, as host cells are responding to multiple morphologies with varying exposure and masking of PAMPs. Additionally, hyphal transition transcriptionally controls *C. albicans* virulence factors, like candidalysin, which also have been shown to modulate immune responses (45–48). Thus, studying live *C. albicans* infection provides a more complex interaction between pathogen and host cell responses which is more representable of *in vivo* interactions.

In addition, *in vivo Candida* infection may not occur in isolation and may occur in the context of a polymicrobial environment, especially for infections of mucosal membranes. Mixed bacterial-fungal infections have been associated with increased disease severity. For example, *in vivo* murine sublethal co-infections of *C. albicans* and bacterial *Staphylococcus aureus* was shown to have 80% mortality, whereas infection of *C. albicans* or *S. aureus* alone resulted in no mortality (49). *C. albicans* is often found together with gram negative *Pseudomonas aeruginosa* (bacterium) in polymicrobial biofilms from cystic fibrosis patients (50,51). Furthermore, Zebrafish models challenged to co-infection with *C. albicans* and *P. aeruginosa* showed reduced survival compared to single infections (52). While polymicrobial infections have been shown to increase risk for host mortality, it is not yet clear how the host cell responds to bacterial and fungal co-infections.

To gain further insight into these interactions, we studied the murine BMDM proteome remodelling after 8-hours of *C. albicans* infection, in combination with the gram-negative bacterial cell wall component LPS or co-infection with the gram-negative bacteria *P. aeruginosa* using Data Independent Acquisition (DIA) based proteomic approaches. Together, we found that while macrophages have a limited measurable response to *C. albicans* infection in isolation, *C. albicans* has targeted protein suppression of important inflammatory macrophage responses to LPS or *P. aeruginosa* infection, including IL-6 and IL-12 secretion.

## Results

### *C. albicans* infected BMDM proteomes consistently regulate a distinct group of inflammatory and anti-inflammatory proteins

*C. albicans* has been shown to induce MAPK and NFκB signalling as well as TNF production in macrophages (35,39). In line with this, we found that infection of murine Bone Marrow Derived Macrophages (BMDMs) with live *C. albicans* stimulated TNF secretion and activation of the p38 MAPK and NFκB pathways (Supplementary Figure 1). To better understand the overall response of BMDMs to *C. albicans* infection, total proteomes were analysed after 8 hours of *C. albicans* infection and compared to unstimulated controls in two independent proteomic experiments, each with 4 biological replicates. The estimated total protein content per cell in unstimulated and *C. albicans* infected BMDMs were not significantly different suggesting the protein content and macrophage size was unchanged during infection (Supplementary Figure 2 A, B). Unstimulated BMDM protein expression from Experiment 1 and 2 were well correlated (Spearman coefficient 0.9774) (Supplementary Figure 2C), although the variation between Experiment 1 and 2 was greater than the variation between the biological replicates within one experiment (Spearman coefficients ranged from 0.98 to 1.0 for comparison of individual replicates within each experiment). This may reflect slight variations in the macrophage differentiation on different days, and for this reason, data from the two experiments was analysed separately. To determine the differential expression of proteins between *C. albicans* infection and unstimulated conditions, regulated proteins were defined as those *with q*-values (Limma calculated adjusted *p*-value) of less than 0.05 and a Log2 fold-change of more than 1 standard deviation from the median. Up-regulated proteins were removed from the list if identified in 2 or less stimulated replicates, while in the down-regulated subset, proteins were removed if identified in 2 or less of the unstimulated replicates. An additional group of proteins were categorised as regulated, where proteins were included in the up-regulated list if proteins were not detected in unstimulated conditions yet present in all stimulated replicates; or down-regulated if present in all replicates of unstimulated group yet not detected in stimulated groups.

In the first experiment, *C. albicans* infected BMDMs up-regulated 162 proteins and down-regulated 108 proteins (Figure 1A). In the second experiment, *C. albicans* infected BMDMs proteome differentially up-regulated 99 proteins and down-regulated 36 proteins (Figure 1B). Analysis of these two experiments showed that the overlap between the regulated protein groups in the two experiments was not perfect (Supplementary Figure 2D and E), especially for Experiment 1 were many of the down-regulated proteins were not replicated in the 2^nd^ experiment. To define a list of proteins robustly regulated by *C. albicans* infection, proteins found up- or down-regulated across the two experiments were collated (Figure 1 C, D; Supplementary Table 1). Within this list of proteins, the fold changes between Experiment 1 and 2 showed a strong positive correlation (spearman correlation r=0.8289) (Figure 1C). The list of up-regulated proteins included several PRRs (Clec4d, Clec4e, TLR2) and signalling proteins that act downstream of PRR activation (IRAK2, CHUK/IKKa) while the NFkB inhibitor NFKBIB/IkBb was down-regulated. In addition, several immune regulatory proteins (ACOD1, IL-1RN and SLAMF7), transcription factors with roles in innate immunity (ATF3, JUNB and CEBPB) and two autophagy associated proteins including UBQLN4 and SQSTM1 (2 isoforms were detected) were also up-regulated on *C. albicans* infection.

**Figure 1.**
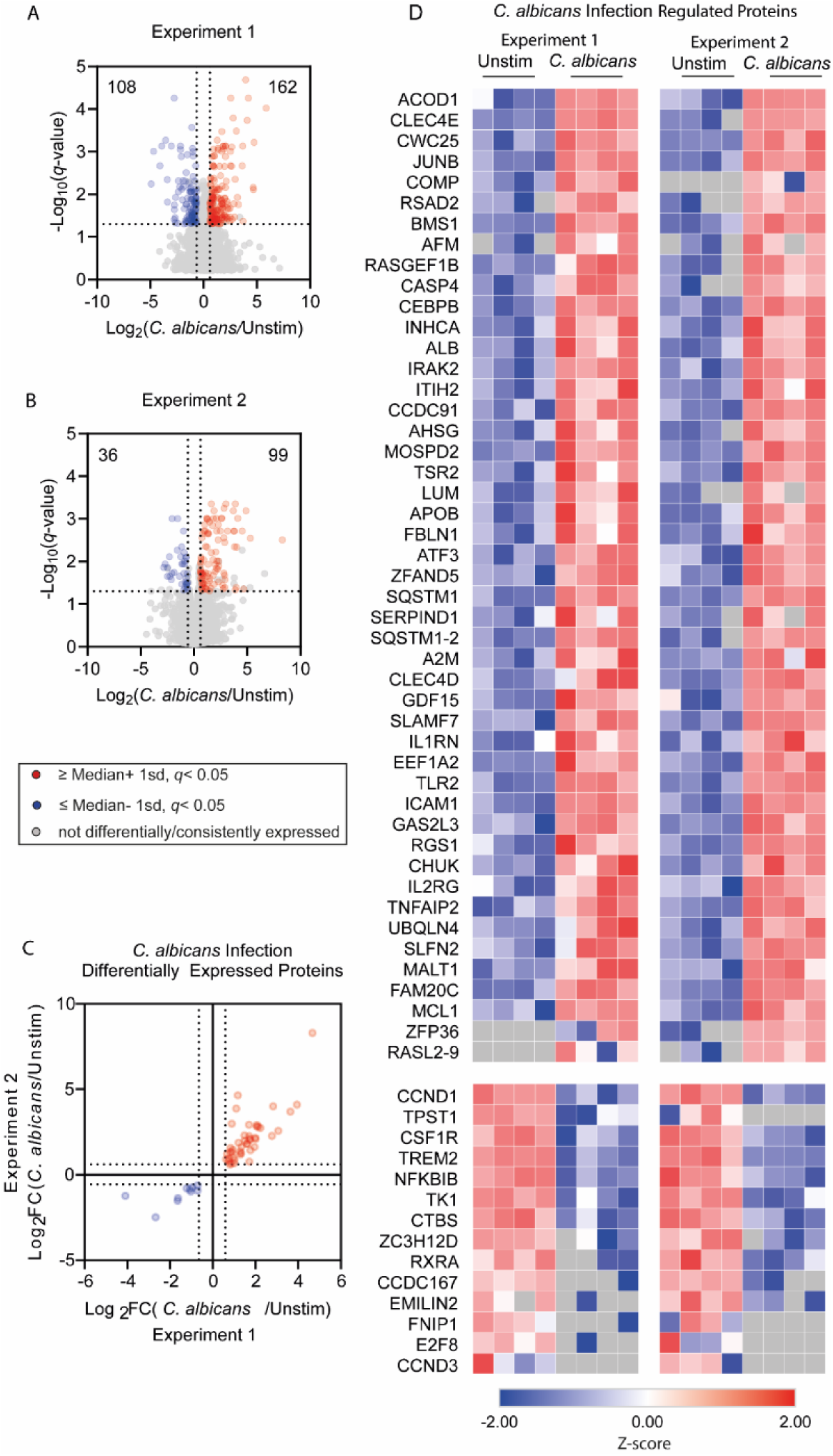
BMDM proteomic response to *C. albicans* infection. (A-D) BMDMs were infected with *C. albicans* for 8 hours and then lysed for proteomic analysis. The experiment was carried out twice on different days, each using 4 biological replicates (separate preparations of macrophages from 4 mice). Differentially expressed proteins were defined as having a fold change of greater than 1 standard deviation away from the median, and a *q*-value <0.05; where up-regulated proteins were present in ≥3 of the infected replicates, and down-regulated proteins were present in ≥3 unstimulated replicates. (A and B) Volcano plots highlight differentially up- and down-regulated proteins as well as non-differentially expressed proteins (red, blue and grey respectively), with fold change cut-offs shown with vertical lines, and *q*-value cut-off as horizontal line. (C and D) A refined subset of proteins which passed the cut-offs for differential expression in both Experiment 1 and 2 was generated. For this subset, an xy plot of log2 fold change values is shown in (C), the Spearman coefficient for this correlation was 0.8289. (D) Shows heat maps for the refined subset of proteins plus an additional group of proteins if proteins were not detected in unstimulated conditions yet present in all replicates in stimulated, or if present in all replicates of unstimulated group yet not detected in stimulated groups. These proteins were plotted in the same order for Experiment 1 and 2 for direct comparison and shown in Supplementary Table 1.

### *C. albicans* infection modulated the LPS response in BMDMs

Overall, the effect of *C. albicans* infection on proteome remodelling was modest, despite the important roles attributed to macrophage infection models *in vivo* (18–20). For this reason, coupled to the finding that *C. albicans* did regulate some proteins with known function in innate immunity, we compared the macrophages response to infection with live *C. albicans* and the changes occurring with LPS or the bacterial pathogen *P. aeruginosa*. Given that *in vivo* exposure to *C. albicans* may occur in a polymicrobial environment, we also examined whether *C. albicans* might modulate the macrophage response to either LPS or *P. aeruginosa* infection.

LPS is a component of gram-negative bacteria outermost membrane and has been used extensively to mimic anti-bacterial responses in cultured macrophages. In terms of proteomic remodelling, both LPS and infection with the gram negative-bacteria *P. aeruginosa* resulted in similar responses after 8 hours in BMDMs (Warner *et al*., manuscript in preparation). We therefore initially used LPS as a simplified model of *C. albicans*-bacterial co-infections. Cytokine secretion was compared in BMDMs infected with *C. albicans*, stimulated with LPS, or co-treated with both. As expected, LPS stimulation induced a robust increase in the secretion of IL-6, IL-12p40, TNF, and IL-10 compared to unstimulated controls. In contrast, *C. albicans* infected BMDMs did not induce a large cytokine response compared to LPS stimulated BMDMs (Figure 2A-D). BMDMs co-treated with *C. albicans* and LPS produced more TNF and IL-10 compared to those treated with LPS alone (Figure 2C, D). Interestingly *C. albicans* significantly suppressed IL-6 and IL-12p40 production in response to LPS (Figure 2A, B). Yet, this suppression was selective as co-treated BMDMs secreted significantly more TNF and IL-10 relative to LPS stimulated BMDMs. The decrease in LPS induced IL-6 and IL-12p40 secretion in co-treated BMDMs was not due to *C. albicans* affecting the viability of the BMDMs during the 8-hour stimulation as there was no significant difference between total number of cells or their survival, as judged by cellular exclusion of DNA binding dyes, between the LPS and co-treated conditions (Supplementary Figure 3).

**Figure 2.**
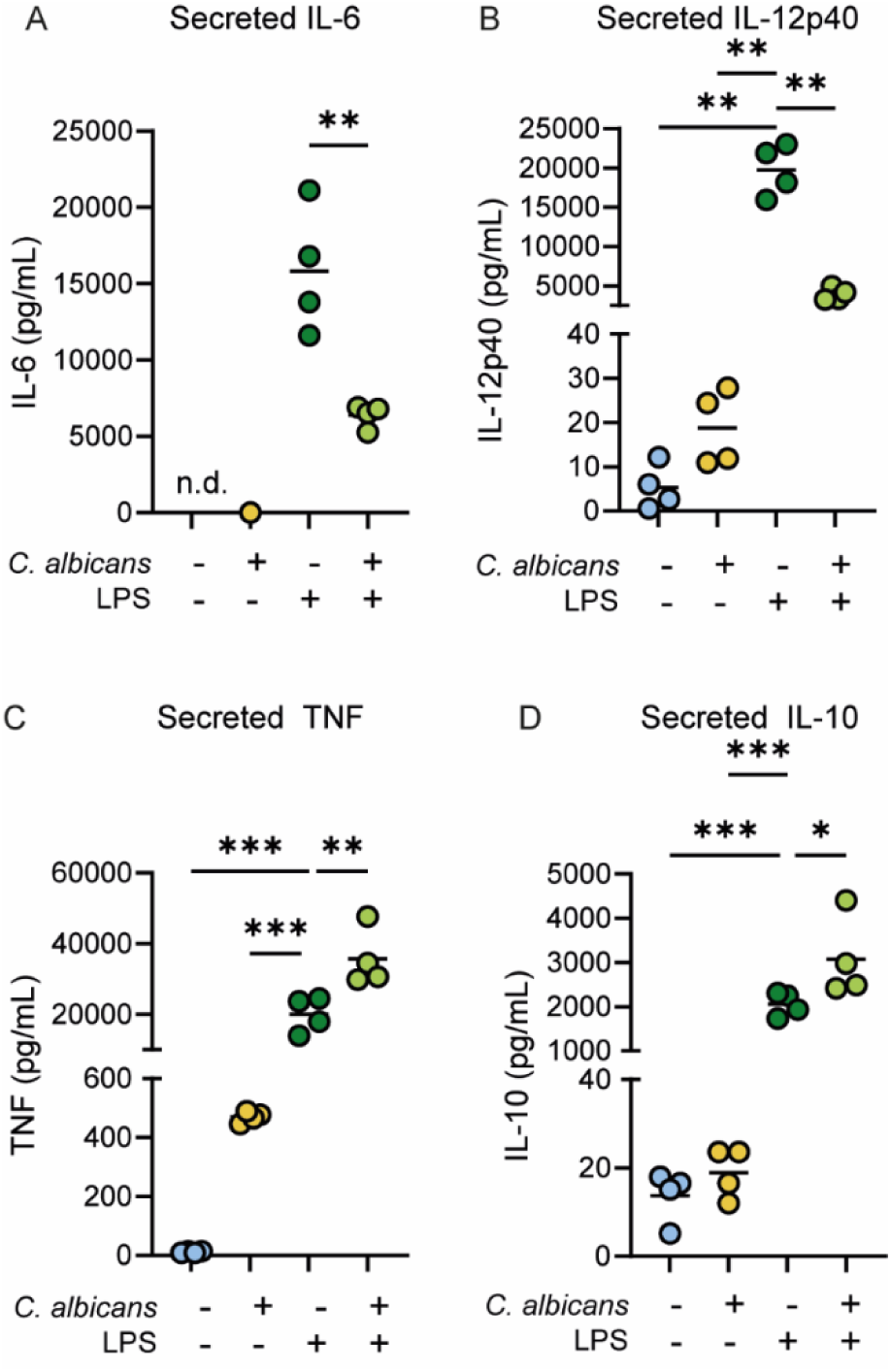
*C. albicans* selectively suppresses LPS induced cytokine production in BMDMs. BMDMs were either infected with live *C. albicans* (MOI 4), stimulated with 100 ng/ml LPS or co-treated with both agents for 8-hours. Culture media was then collected and analysed for the levels of IL-6 (A), IL-12p40 (B), TNF (C) or IL-10 (D). Data shows 4 biological replicates for the macrophages with individual replicates shown by symbols and mean values by a line. Significant differences were determined by Students ttest (A) or one-way ANOVA with Turkeys multiple comparison tests (C, F=50.56, p<0.0001; D, F=40.3, p<0.0001) or Welch’s ANOVA with Dunnett’s multiple comparison tests (B, F=66.43, p<0.0001). For compassions to the LPS alone condition, adj. p < 0.05 is indicated by * , < 0.01 by ** and < 0.001 by ***.

To determine whether the effect of *C. albicans* on LPS cytokine secretion was due to upstream suppression of known TLR4 signalling (38), these pathways were examined at 0.5 and 2.5 hours of treatment (Supplementary Figure 4). In line with previous reports, LPS induced the activation of the p38 and ERK1/2 MAPK cascades, as judged by phosphorylation of their TXY activation motifs, or the phosphorylation of CREB, a known downstream target of MAPK signalling in macrophages (53). IKK signalling was also activated, as determined by the phosphorylation of the p105 NFkB subunit. *C. albicans* infection with *C. albicans* alone only weakly activated signalling compared to LPS. Co-treatment with LPS and *C. albicans* resulted in similar levels of MAPK and NFkB activation (Supplementary Figure 4), suggesting *C. albicans* did not affect IL-6 and IL-12p40 production by modulating the initial signalling induced by TLR4.

To examine if the suppression of IL-6 by *C. albicans* was specific for its induction by TLR4 activation, the effect of *C. albicans* on IL-6 production in response to TLR agonists was examined. In line with the results for LPS, co-treatment with *C. albicans* and either Pam3CSK (TLR1/2), R848 (TLR7/8), CL097 (TLR7/8) or CpG DNA (TLR9) resulted in lower IL-6 production relative to the TLR agonist alone (Figure 3A). *C. albicans* has been reported to activate the C-type lectin receptor Dectin-1 with β-glucan structures in the fungal cell wall (40,54). Unlike live *C. albicans*, depleted zymosan, a selective Dectin-1 agonist, was able to induce IL-6 production in macrophages, although not as strongly as LPS (Figure 3B). LPS co-treatment with depleted zymosan did not reduce IL-6 production compared to LPS stimulation alone (Figure 3B), suggesting that Dectin-1 activation by *C. albicans* was not responsible for the effects seen on LPS induced IL-6 production seen in Figure 2A. In line with the suppression being independent of Dectin-1, *C. albicans* infection also suppressed depleted zymosan induction of IL-6 (Figure 3C).

**Figure 3.**
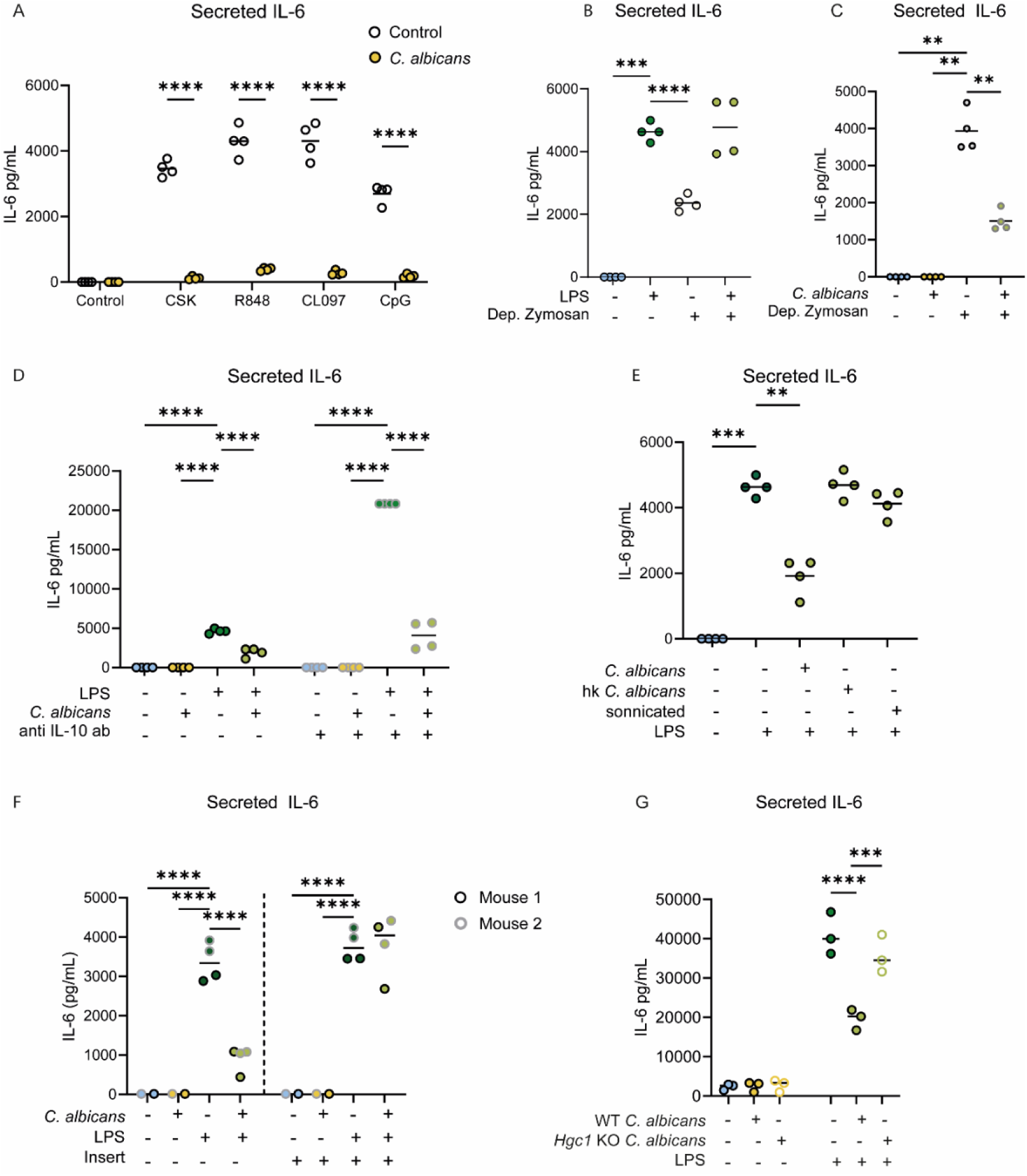
Suppression of LPS production in BMDMs by *C. albicans* requires direct contact with live *C. albicans*. To investigate the effect of *C. albicans* on IL-6 production, macrophages were treated in the described ways and then IL-6 levels were determined in the culture media. (A) BMDMs were treated for 8h with 100 ng/ml Pam3CSK, 250 ng/ml R848, 1 µg/ml CL097 or 2µM CpG in the presence or absence of infection with *C. albicans* (MOI 4). (B, C) BMDMs were treated with the indicated combinations of 100 ng/ml LPS and 200 µg/ml depleted zymosan or infection with *C. albicans* (MOI 4) for 8h. (D) BMDMs were pre-treated with an IL-10 neutralising antibody 15-minutes prior to treatment for 8h with the indicated combinations of *C. albicans* (MOI4) and 100 ng/ml LPS. (E) BMDMs were treated with 100 ng/ml LPS and either live *C. albicans* (MOI4) or equivalent amounts of heat-killed *C. albicans* or sonicated *C. albicans*. (F) BMDMs were seeded in Transwell® plates. *C. albicans* was either inserted into Transwell insert or directly into the wells with the macrophages. LPS was added directly onto BMDMs regardless of insert presence. (G) BMDMs were co-treated with LPS and either wild type (WT) *C. albicans* or *Hgc1* knockout (KO) *C. albicans*. Graphs show results from 3 or 4 biological replicates except (F) were 2 technical replicates (separate culture wells) on 2 biological replicates are shown. Data was analysed by two way ANOVA with Sidak’s multiple comparison tests (A, D, F, G) or Welch’s ANOVA (B, C, E) with Dunnett’s T3 tests F and P values for the ANOVA are given in Supplementary Table 2. For compassions to the LPS alone condition (B, D-E) or to the appropriate PAMP (A, C), adj. p < 0.05 is indicated by *, < 0.01 by **, <0.001 by *** and < 0.0001 by ****.

In response to TLR stimulation, macrophages secrete IL-10 which can activate a feedback loop that inhibits IL-6 and IL-12p40 secretion. As IL-10 was elevated in LPS and *C. albicans* co-treated BMDMs relative to LPS (Figure 2D), a potential role for IL-10 in the suppressive effect of *C. albicans* on TLR induced IL-6 production was tested using an IL-10 neutralising antibody. As expected, treatment of the BMDMs with the IL-10 antibody increased LPS stimulated IL-6 production (Figure 3D). *C. albicans* was, however, still able to inhibit LPS induced IL-6 secretion in the presence of the IL-10 antibody suggesting the effect of *C. albicans* is independent of IL-10 (Figure 3D). To determine whether *C. albicans* needed to be alive to suppresses the LPS induction of IL-6, macrophages were treated with either heat-killed or sonicated *C. albicans* in combination with LPS. This showed that only live *C. albicans* suppressed the LPS induced IL-6 response (Figure 3E).

It is possible that *C. albicans* could modulate IL-6 production through secreted factors such as toxins, proteases or quorum sensing microbial communication molecules. To determine whether IL-6 suppression requires direct contact between the macrophage and *C. albicans*, 0.4 μm pore polycarbonate cell culture inserts were used to physically separate *C. albicans* from BMDMs, while still allowing interaction with secreted molecules and LPS. In conditions without inserts (allowing physical interactions) *C. albicans* suppressed IL-6 as previously shown. Interestingly, separation of *C. albicans* from BMDM — through use of the insert — prevented the suppression of IL-6 secretion in co-treated BMDMs (Figure 3F). Thus, contact between *C. albicans* and BMDMs appears to be necessary for selective suppression of LPS induced IL-6.

*C. albicans* is a polymorphic pathogen, meaning it changes from yeast to hyphae depending on environmental queues. Hyphal formation is triggered during *in vitro* infections, when *C. albicans* is in the presence of FBS supplemented media, and when deprived of nutrients within the macrophage phagosome. The hyphae G1 cyclin-related gene *(HGC1)* is required for morphogenesis of yeast to hyphae transition; thus, *Hgc1* KO (knockout) strains are locked in the yeast morphology *C. albicans* (55). To determine whether hyphal transition may affect IL-6 suppression in BMDMs, wild type *C. albicans* and *Hgc1 knockout* strains were compared for their ability to affect IL-6 production. Interestingly, induction of IL-6 by LPS was not significanly suppressed by *Hgc1* KO co-treatment, although there was a trend towards suppression (Figure 3G). Taken altogether, this data suggests *C. albicans* suppression of IL-6 production in response to TLR/CTLR activation, is independent of IL-10, but dependent on *C. albicans* being alive and having a polymorphic phenotype.

### *C. albicans* broadly suppresses LPS up-regulated proteome

To further examine the effect of *C. albicans* on the macrophages LPS response, proteomics was used to identify LPS induced changes in macrophages in the presence or absence of live *C. albicans*. Initially, we focused on the subset of genes identified in Figure 1 as being regulated by *C. albicans* infection, comparing single stimulation of LPS or *C. albicans* infection. The proteins down-regulated by *C. albicans* were more strongly down-regulated by LPS (Figure 4). The proteins up-regulated by *C. albicans* infection could be divided into 2 main categories – those that were also regulated by LPS and those that were more selective for *C. albicans* infection (Figure 4). Proteins regulated by both treatments generally showed a higher up-regulation with LPS than with *C. albicans* infection. This group contained multiple proteins, such as JUNB, IL1RN, ICAM1, TNFAIP2, SLFN2, TLR2 and ATF3 which are all known to be regulated by MAPK or NFκB (56–63). Their increased up-regulation by LPS is consistent with the finding that LPS is a stronger activator of these pathways than infection with *C. albicans* (Supplementary Figure 4).

**Figure 4.**
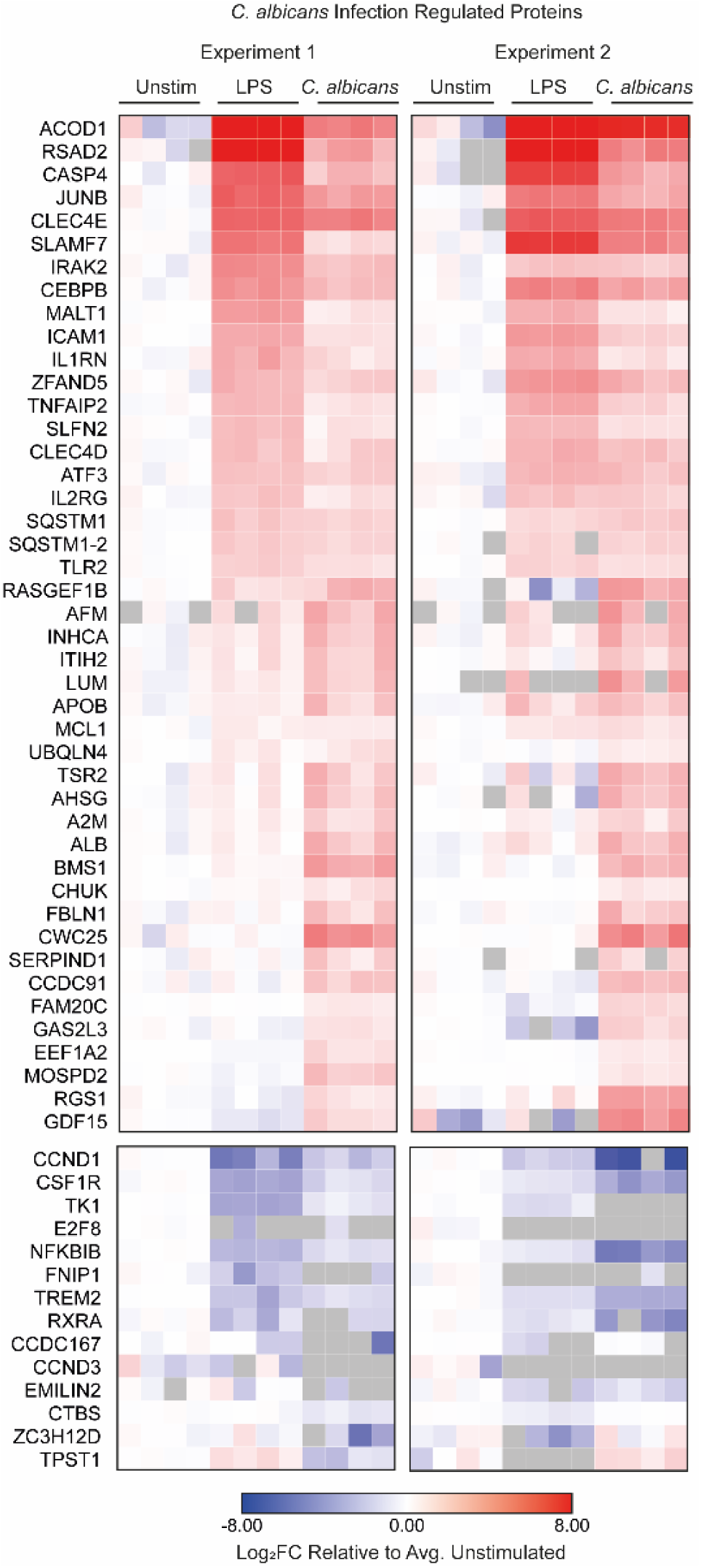
Comparing protein expression of LPS stimulated BMDMs of a *C. albicans* infection regulated protein subset. The heatmap shows log2 fold change (relative to the average on the untreated BMDM replicates) for the refined list of proteins that were down-regulated or up-regulated following *C. albicans* infection from proteomic Experiment 1 and 2 (as defined in Figure 1). Data is shown for the unstimulated BMDM, 8 hour LPS stimulation and 8 hour *C. albicans* infection conditions. *C. albicans* up-regulated proteins which were not present in the unstimulated conditions were not included in this heatmaps as the data plotted is relative to the average unstimulated concentrations (Zfp36, Rasl2-9).

Analysis of the LPS response across the 2 proteomic experiments resulted in the identification of 2 groups of consistently regulated proteins, a group of 84 down-regulated and 349 up-regulated protein (according to the specifications outlined in the methods Supplementary and Figure 5). To look at the effect of co-treatment, the Log2 fold change for the *C. albicans* and LPS group relative to the LPS group alone was calculated. Analysis of these values in the LPS up- or down-regulated protein groups showed that the median fold change between co-treatment and LPS alone was strikingly lower for LPS up-regulated proteins than proteins either down-regulated or unaffected by LPS (Figure 5A, B). A more detailed analysis of LPS down-regulated proteins showed no clear effects of co-treatment with *C. albicans* infection across this group of proteins (Figure 5 C, D). In contrast, *C. albicans* infection appeared to show a consistent suppression of the BMDM LPS up-regulated protein subset. This subset showed the majority of the protein expression in co-stimulated BMDMs was less than the expression in LPS stimulated BMDMs; as shown by fold changes plotted less than 0 in the volcano plots (Figure 5 E, F) with a similar pattern between the two experiments (Figure 5G).

**Figure 5.**
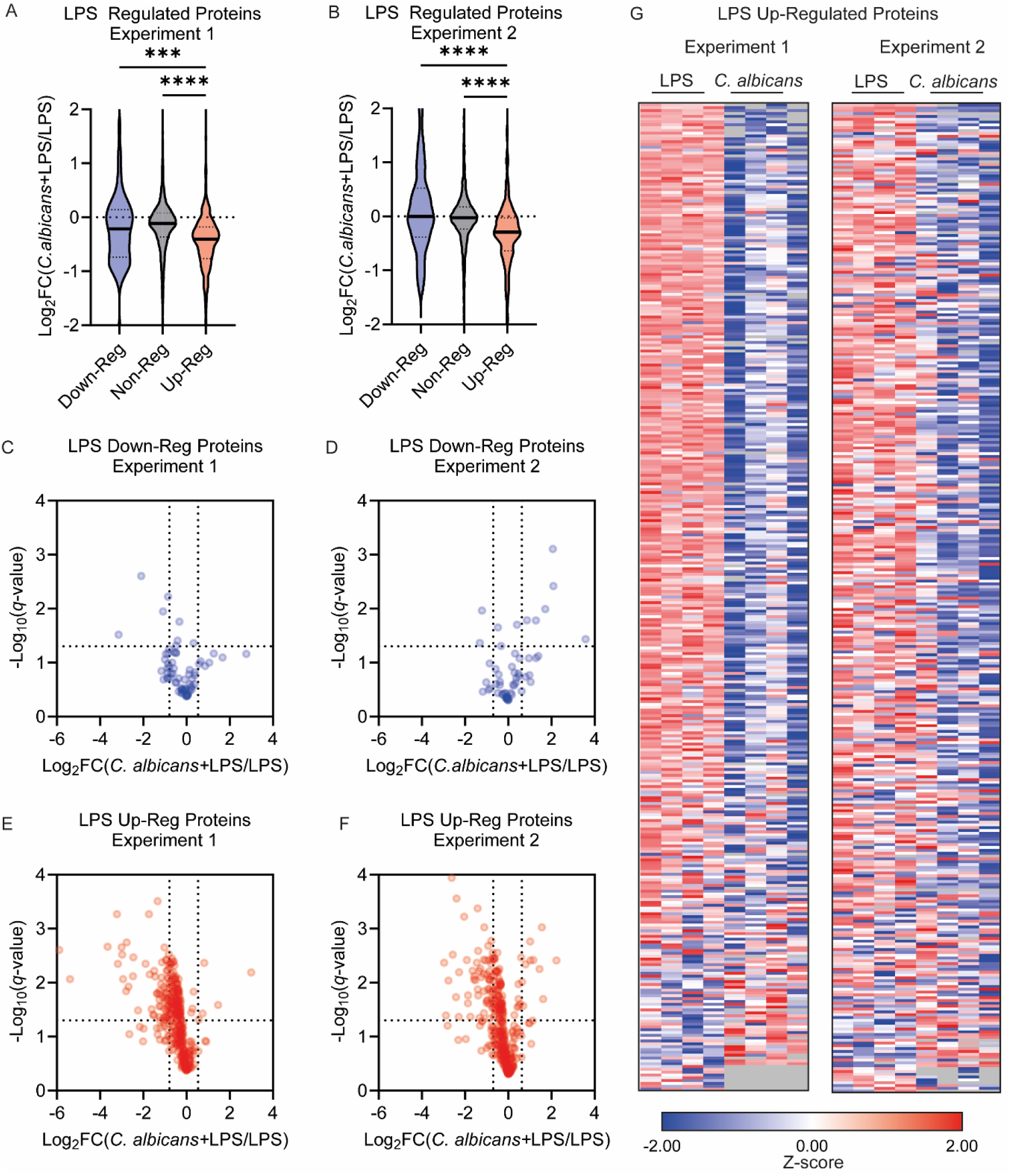
*C. albicans* selectively suppresses a subset of LPS up-regulated in BMDMs. BMDMs were stimulated with 100ng/ml LPS for 8 hours in the presence or absence of infection with *C. albicans*, and then lysed for proteomic analysis. Results show data from 2 independent experiments each with 4 biological replicates. Based on the analysis in Supplemental Figure 5, proteins were divided into those showing a robust up or down regulation by LPS relative to unstimulated cells and those showing no consistent regulation (non-reg). The effect of treatment with *C. albicans* and LPS relative to LPS for proteins in these 3 groups was then calculated. Violin plots of log2 fold changes for the LPS + *C. albicans* condition relative to LPS alone for the 3 groups of LPS regulated proteins are shown in (A) and (B). Volcano plots illustrating the effect of *C. albicans* on the LPS down-regulated proteins are shown (C) and (D) and for up-regulated proteins in (E) and (F). (G) The heatmap showing the effect of *C. albicans* on the subset of proteins that were upregulated by LPS. The heatmap was ordered by the degree of repression in the presence of *C. albicans* seen in Experiment 1. Full list of proteins for the heatmap is given in (Supplementary Table 3). (A) and (B) were analysed by Kruskal-Wallis test (p<0.0001 for both A and B) Dunn’s multiple comparison tests. p < 0.001 is indicated by *** and p < 0.0001 by ****.

Given these global effects, the proteomic data was further mined to look at the effect of *C. albicans* on LPS stimulated BMDM expression of proteins known to be important in the macrophage response. The data for individual proteins in Figure 6 is from Experiment 2; however, the same results were obtained in Experiment 1 and the corresponding data is shown in Supplementary Figure 6. Consistent with the secreted cytokine data, the induction of IL-12p40 by LPS was suppressed by *C. albicans* infection (Figure 6A), while TNF was increased in co-treated BMDMs compared to LPS stimulated BMDMs (Figure 6B). Identification of IL-6 and IL-10 were insufficient to report meaningful results, as they were not consistently detected in the proteomic dataset. Based on the proteomic data, intracellular levels of IL-1β and IL-18 were increased by LPS treatment and in both cases this induction was reduced when the macrophages were also infected with *C. albicans* (Figure 6C and D). IL-27 and IL-1RN were also detected in the proteomics and induced by LPS; however, these cytokines expression were not supressed by *C. albicans* infection (Figure 6E, F). Similar to what was observed with cytokines, the induction of some chemokines by LPS was suppressed by co-treatment with *C. albicans*, as in the case of CCL3, CCL5 and CXCL10, while CCL2, CCL4, CCL7 and CXCL16 were unaffected (Figure 6G-N). CXCL2 was unusual in that it showed a synergistic induction by a combination of LPS and *C. albicans* (Figure 6L). An important component of the macrophages response to inflammatory stimuli is the production of prostaglandins, whose synthesis is controlled by the level of Ptgs2 (COX-2). Ptgs2 was induced by LPS; however, this was not reduced by co-treatment with *C. albicans* infection (Figure 6O). Interestingly, LPS induction of Nitric oxide synthase (NOS2), an important macrophage anti-microbial protein, was suppressed by co-treatment with *C. albicans* (Figure 6P). Additionally, anti-inflammatory protein, ACOD1, which is significantly regulated by *C. albicans* and LPS separately, was suppressed in co-treated BMDMs relative to LPS stimulation alone (Figure 6 Q and Supplementary Figure 6 Q). The transcription factors highlighted here (ATF3, JUNB and CEBPB) were induced by *C. albicans* infection alone, and either significantly increased during co-infection or not significantly different to LPS simulation alone (Figure 6 R-T).

**Figure 6.**
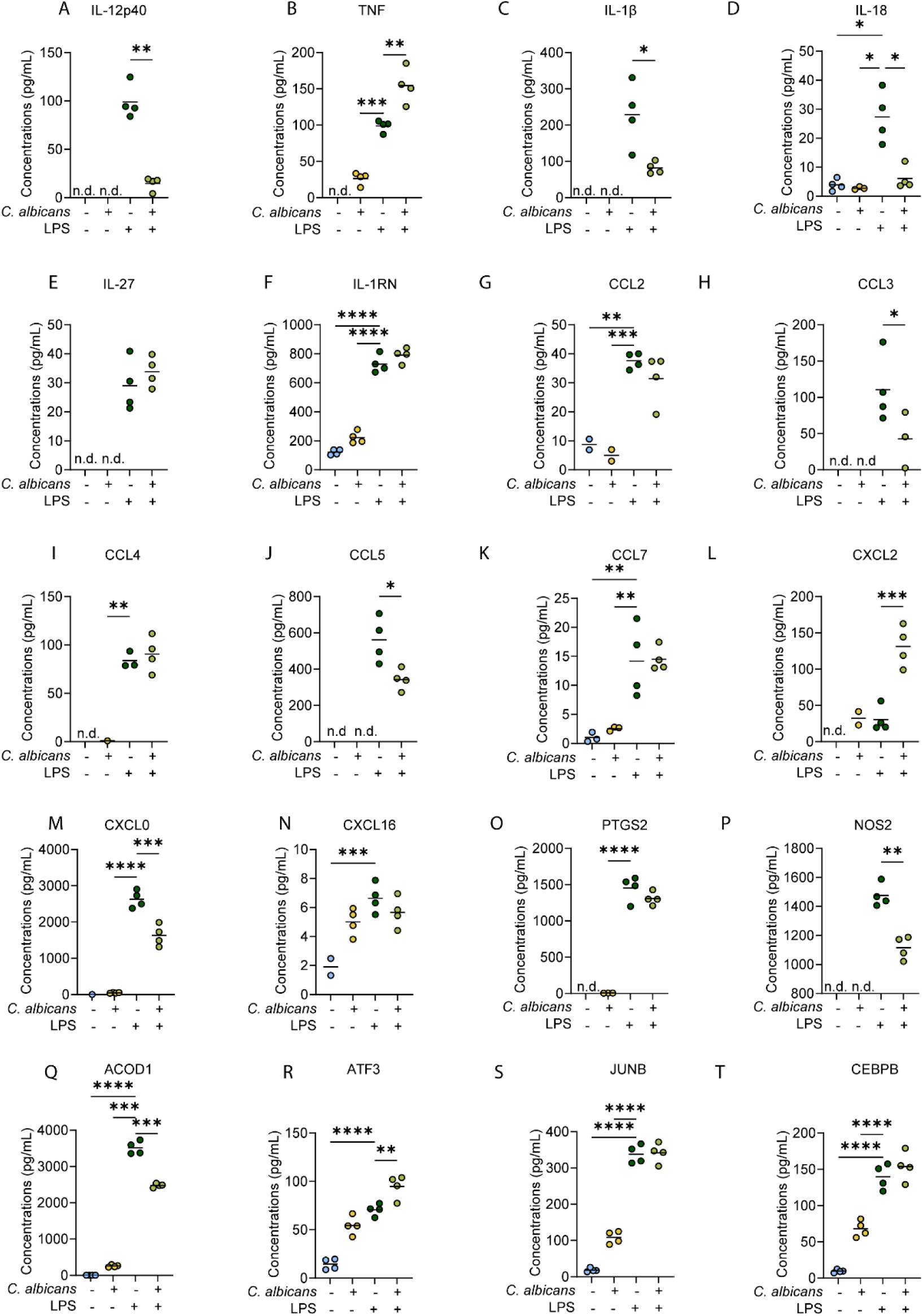
Effect of *C. albicans* on the induction of selected LPS regulated proteins in BMBMs. (A-T) The levels of the indicated proteins in response to *C. albicans* infection and/or LPS stimulation were derived from the proteomic dataset described in Figure 5. Graphs show values for 4 biological replicates with mean values indicated by lines. Data from one experiment is shown. Similar results were obtained from the 2^nd^ proteomic dataset and are shown in Supplementary Figure 6. Data was analysed by Students ttest, one-way ANOVA or Welch’s ANOVA as appropriate and statistics are summarised in Supplementary Table 4. All post-hoc comparisons were made relative to the LPS alone condition. An adjusted *p*-value of < 0.05 is indicated by *, < 0.01 by ** , < 0.001 by ***,< 0.0001 by ****.

### *C. albicans* modulates the *P. aeruginosa* response in BMDMs

As LPS is a component of gram-negative bacterial outer membrane, LPS is a common agonist used to model bacterial infection, such as *Pseudonomas aeruginosa*. To determine whether the suppressive capabilities of *C. albicans* on LPS stimulated BMDMs could be replicated in a live bacterial-fungal co-infection, BMDMs were co-infected with *C. albicans* and *P. aeruginosa* for 8 hours followed by media collection for secreted cytokine analysis and lysis of BMDMs for proteomic analysis. Consistent with what was shown for *C. albicans* co-treated responses with LPS, *C. albicans* selectively suppressed *P. aeruginosa* induction of IL-6 and IL-12p40 in co-infected conditions relative to *P. aeruginosa* infection alone, while TNF and IL-10 levels escaped suppression (Figure 7A-D).

**Figure 7.**
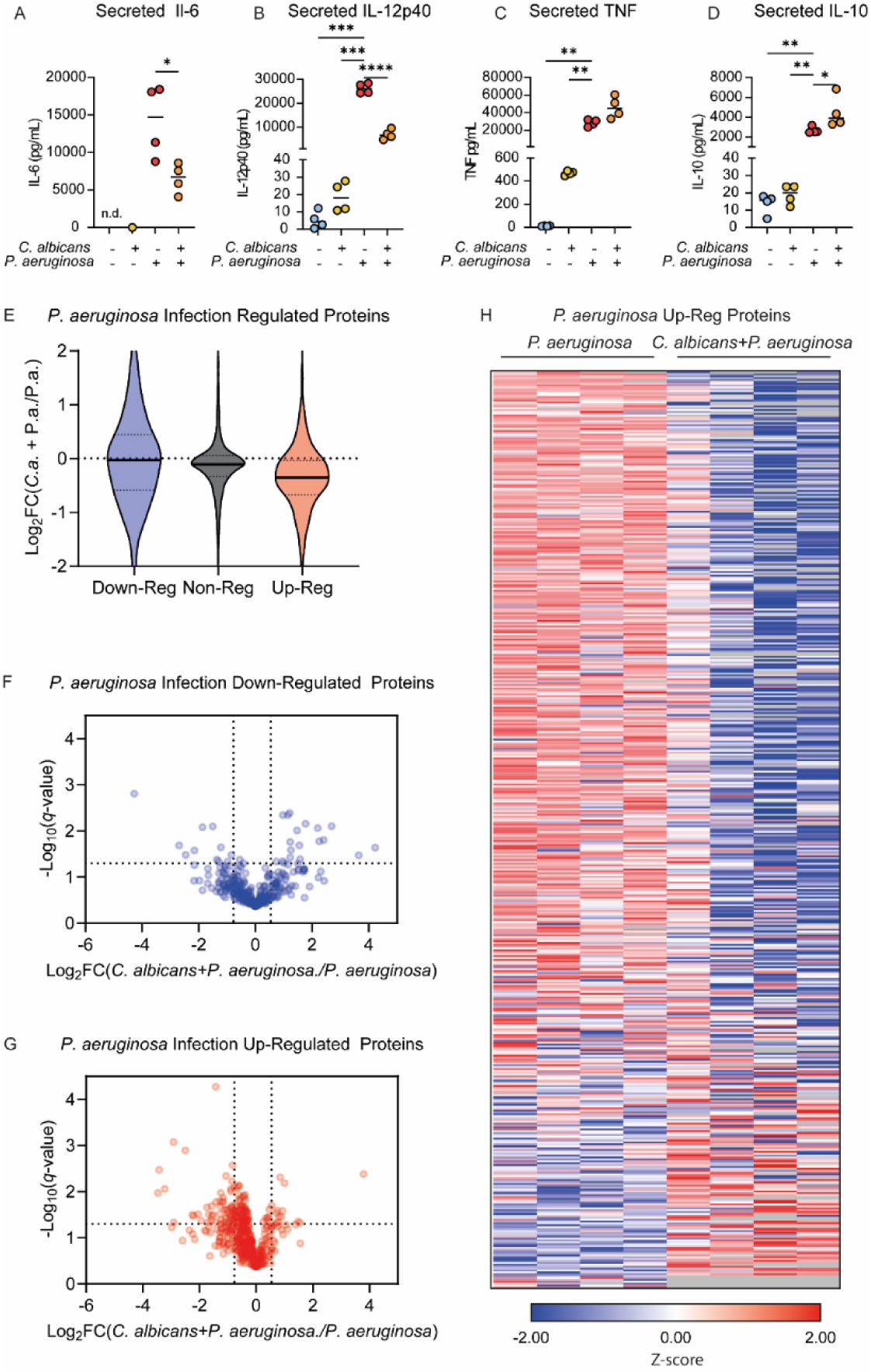
*C. albicans* suppresses the BMDMs response to *P. aeruginosa* infection. BMDMs were left un-treated or infected with *C. albicans* (MOI4), *P. aeruginosa* (MOI 10) or co-infected with both organisms for 8-hours. Media was collected for subsequent analysis of secreted cytokine analysis and the cells were lysed for proteomic analysis; 4 biological replicates were analysed per condition (A-D). The levels of IL-6 (A), IL-12p40 (B), TNF (C) and IL-10 (C) present in the media at 8h are shown. Graphs show individual biological replicates as symbols with the means indicated by a line. (E-H) Using the analysis described in Supplementary Figure 7, proteins present in the proteomic dataset were divided into those up-regulated by *P. aeruginosa* infection relative to un-infected cells, those that were downregulated and those that did not show a strong regulation (non-reg). The effect of co-infection with *C. albicans* relative to infection with just *P. aeruginosa* for proteins in these 3 groups was then calculated. (E) The violin plot shows log2 fold changes for the *P. aeruginosa* and *C. albicans* co-infected condition relative to the *P. aeruginosa* infection alone for the groups of the *P. aeruginosa* regulated proteins (F, G) Volcano plots illustrating the effect of co-infection with *C. albicans* the on proteins that were down-regulated (F) or up-regulated (G) by *P. aeruginosa* infection in BMDMs. (H) Heatmap showing the effect of *C. albicans* co-infection on the subset of proteins that were up-regulated by *P. aeruginosa* infection. Heatmap was ordered by the degree of repression in the presence of *C. albicans*. Full list of proteins for the heatmap is given in (Supplementary Table 5). Statistical testes used were (A) Students ttest, (B) Welch’s ANOVA, F=179.4 p < 0.0001, (C) one way ANOVA, F=27.42 p < 0.0001 (D) Welch’s ANOVA F=703.8, p < 0.0001 and (E) Kruskal-Wallis test p < 0.0001. Dunnett’s T3 multiple comparison tests were used for (B) and (D), Tukey’s tests for (C) and (D) and Dunn’s tests for (E). For comparisons to the *P. aeruginosa* infection condition in (A-D) an adj. *p* < 0.05 is indicated by *, < 0.01 by ** and < 0.01 by ***, and < 0.001 by ****.

Proteomic analysis of *P. aeruginosa* infection response in BMDMs identified 385 down-regulated and 408 up-regulated (based on the criteria described in the methods and Supplementary Figure 7). In addition, 79 proteins were included in the up-regulated subset which were detected in all of the *P. aeruginosa* infected BMDMs replicates but none of the untreated ones. To determine how *C. albicans* may modulate the *P. aeruginosa* infection response in BMDMs, the log2 FC for *P. aeruginosa* infection versus co-infection was determined for proteins identified as regulated by *P. aeruginosa* infection (Figure 7E). The median Log2 fold change between co-infected and *P. aeruginosa* infection was lower for *P. aeruginosa* up-regulated proteins, compared to the down-regulated or non-regulated protein subsets — consistent with the results for *C. albicans* and LPS shown in Figure 5. BMDM protein expression for co-infected relative to single *P. aeruginosa* infections for the *P. aeruginosa* down-regulated and up-regulated protein subsets was visualised by volcano plots. Similar to the results for LPS, these did not show a consistent effect of *C. albicans* on proteins down-regulated by *P. aeruginosa* infection (Figure 7F). For proteins up-regulated by *P. aeruginosa* infection, there was a clear trend for expression levels to be reduced by co-infection with *C. albicans* (Figure 7G, H). While the strength of suppression on protein expression levels was varied, the overall trend of suppression appeared to target the majority of up-regulated proteins. Intriguingly, several important antimicrobial proteins were in this group of suppressed proteins.

As for the experiments with LPS, the effect of co-infection with *C. albicans* and *P. aeruginosa* was examined for the same set of proteins with known immune function (Figure 8A-T). Mirroring the results for LPS, the induction of IL-12p40, IL-18, CCL5, by *P. aeruginosa* infection was reduced by co-infection; while IL-1β, CCL3 and CXCL10 expression trended towards suppression in co-infected BMDMs (Figure 8A,C,D,H,J,M). Also similar to LPS, CXCL2 induction was more strongly induced by co-infection than either single infection (Figure 8L). TNF, IL-27, IL-1RN, CCL2 and CXCL16 were not effected by *C. albicans* co-infection relative to *P. aeruginosa* infection alone (Figure 8B,E,F,G,N). CCL4 and CCL7 showed reduced levels in co-infected relative to *P. aeruginosa* infected BMDMs, which is in contrast to the results for LPS where they were not affected (compare Figure 8I and K to 6I and K). PTGS2 was not affected by co-infection (Figure 8O) while NOS2 was suppressed during *C. albicans* co-infection compared to *P. aeruginosa* infection (Figure 8P), again mirroring the results obtained in the experiments with LPS (Figure 6O, P). Interestingly, anti-inflammatory ACOD1 was also suppressed during co-infection relative to *P. aeruginosa* infection (Figure 8Q). Additionally, several transcription factors, ATF3, JUNB and CEBPB which were highlighted to be induced by *C. albicans* infection appeared to be significantly or trended towards induction in co-infected conditions relative to *P. aeruginosa* infection (Figure 8R-T).

**Figure 8.**
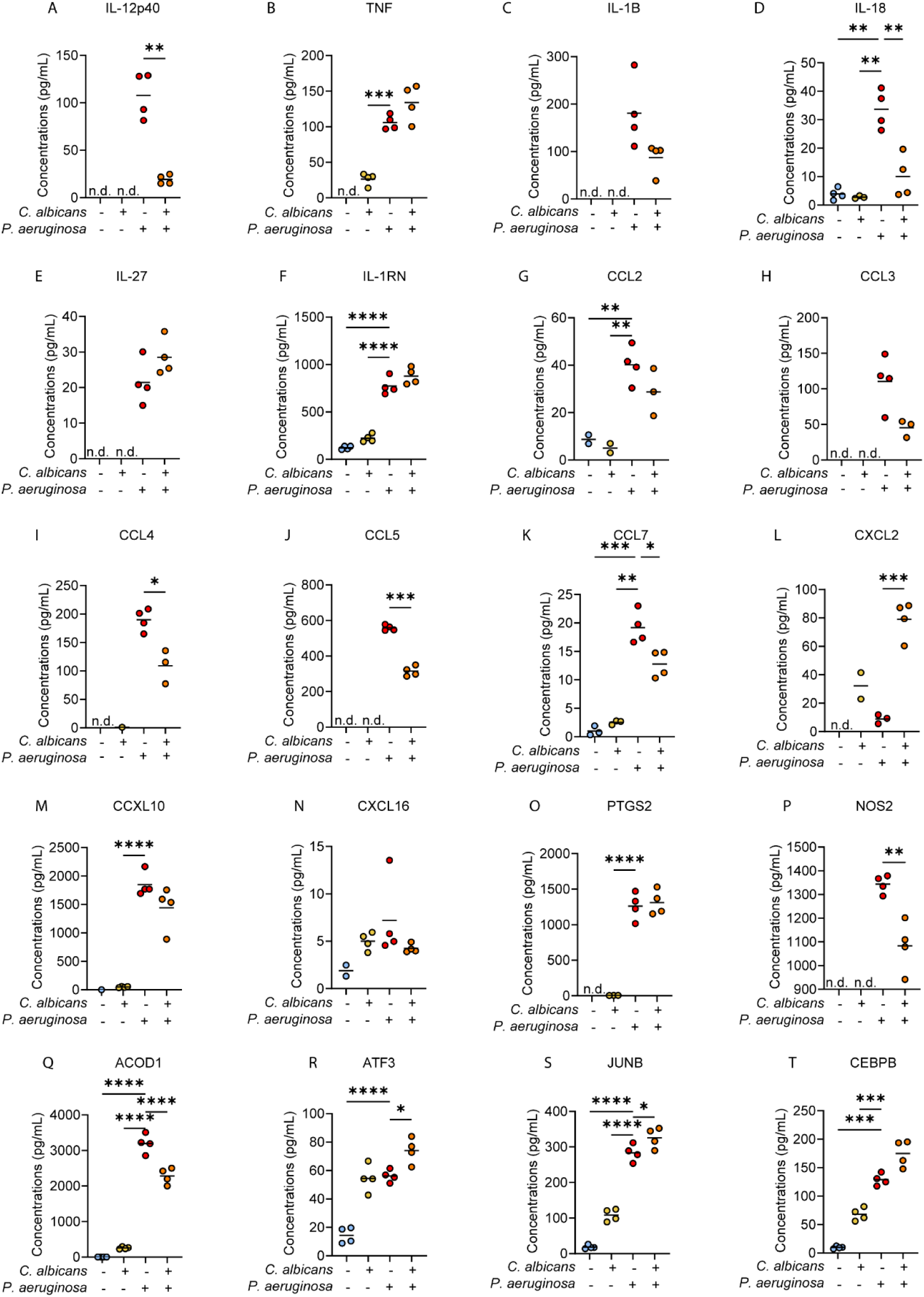
Effect of *C. albicans* on the induction of selected proteins regulated by *P. aeruginosa* infection in BMBMs. (A-T) The levels of the indicated proteins in response to *C. albicans* infection, *P. aeruginosa* infection or co-infection were derived from the proteomic dataset described in Figure 7. The graphs show values for 4 biological replicates with mean values indicated by lines. Data was analysed by Students ttest, one-way ANOVA or Welch’s ANOVA as appropriate and statistics are summarised in Supplementary Table 4. All post-hoc comparisons were made relative to the LPS condition. An adjusted *p*-value of < 0.05 is indicated by *, < 0.01 by ** , < 0.001 by *** and < 0.0001 by ****.

Overal, we have identified an interesting and potentially physiologically relevant phenotype where *C. albicans* appears to selectivley suppress LPS and *P. aeruginosa* induced secreted inflammatory mediators like IL-6 and IL-12, as well as broadly suppress LPS and *P. aeruginosa* up-regulated intracellular protein expression. While the mechanism was not identified, this work provides a comprehensive characterisation of the BMDM responses to *C. albicans* infection, which can support future work.

## Discussion

The results presented in this paper suggest that despite the presence of multiple PAMPs in its cell wall, *C. albicans,* is able to substantially avoid directly activating macrophages. This could indicate that many of the PAMPs in the cell wall are hidden and not directly available for activation by cell surface receptors on the macrophages. *C. albicans* infection of BMDMs resulted in a much more muted MAPK and NFκB signalling response, cytokine induction and proteomic changes compared to LPS or live *P. aeruginosa* infection. This was unexpected, given that *C. albicans* can induce a strong inflammatory response *in vivo* and that macrophage depletion *in vivo* results in significantly increased fungal burdens and decreased survival in invasive candidiasis models (18–20). One potential explanation for this difference would be that opsonisation of the *C. albicans*, either by antibodies or complement, occurs *in vivo*, and that this is important for fully activating the macrophages.

The response of macrophages to *C. albicans* infection was found to be much weaker than that to the TLR4 agonist LPS or infection with *P. aeruginosa.* While this is consistent with *C. albicans* being less able to activate PRR induced signalling, we also unexpectedly found that *C. albicans* infection was able to repress components of the macrophage response to LPS or *P. aeruginosa* infection. This included both IL-6 and IL-12p40 production – two cytokines known to be involved in control *Candida* infection *in vivo* (64,65). The production of nitric oxide is another function of macrophages that has been linked to controlling *Candida* infection. The production of nitric oxide in macrophages requires the enzyme NOS2, which is induced in macrophages of TLR stimulation or bacterial infection (66). In this study infection of macrophages with *C. albicans* did not induce measurable NOS2 expression. In contrast, *P. aeruginosa* infection or LPS treatment robustly induced NOS2 protein expression, and this was reduced by co-treatment with *C. albicans*, similar to what was observed for IL-6 or IL-12p40 (Figure 6 and 8). The ability of *C. albicans* to reduce LPS induced nitric oxide production has been observed previously. Interestingly, this was found to require live *C. albicans* and direct contact of the *C. albicans* and the macrophage (67), similar to what was observed for suppression of IL-6 here.

While *C. albicans* only induced a limited proteome remodelling response, this study did highlight a core set of proteins consistently regulated by *C. albicans* infection across two independent experiments, and it is possible some of these proteins may affect the ability of macrophage to respond to other signals. One protein that may be of note in this respect is the transcription factor ATF3. In macrophages ATF3 has been described as a negative regulator of the inflammatory response, with loss of ATF3 function resulting in elevated induction of IL-6, IL-12 and NOS2 is response to LPS (68–70). In the proteomics experiments reported here, ATF3 was induced by both *C. albicans* infection or LPS stimulation but was one of the few proteins to show a higher induction when the macrophages were co-treated with LPS and *C. albicans* compared to LPS alone. This elevated ATF3 would be consistent with the reduction in IL-6, IL-12p40 and NOS2 expression in the co-treated cells relative to LPS alone (Figure 6). Similar results were obtained in the experiments looking at *C. albicans* and *P. aeruginosa* co-infection (Figure 8). ATF3 is a bZip transcription factor and has been reported to inhibit transcription when it acts as dimer. Alternatively it may form heterodimers with members of the AP-1 subfamily of bZip transcription factors such as ATF2, c-Jun, JunB and JunD and in these cases it may promote transcription (reviewed in (71)). Interestingly, JunB was also induced by LPS stimulation or *C. albicans* infection; however, this transcription factor would be less likely to mediate suppression of LPS induced proteins by *Candida* was work with JunB knockout macrophages indicates that it promotes rather than inhibits IL-12p40 production (72).

ACOD1 was up-regulated during *C. albicans* infection (Figure 1). During inflammatory macrophage activation, ACOD1 diverts cis-aconitate from the Citric Acid Cycle, producing itaconate, which has been found to limit inflammatory responses in macrophages (73–76). ACOD1 was up-regulated more strongly by *P. aeruginosa* infection or LPS compared to *C. albicans* infection. Intriguingly, despite being up-regulated by *Candida* infection on its own, ACOD1 was induced less strongly in macrophages co-treated with *C. albicans* and LPS compared to LPS alone. As such, while ACOD1 may have an anti-inflammatory function it is unlikely to be mediating the repressive effect of *C. albicans* in LPS responses.

Perhaps the most intriguing result presented here, was the broad impact on suppressing BMDM protein expression in LPS stimulation or *P. aeruginosa* infection. This was described both by the broad suppression of LPS or *P. aeruginosa* up-regulated proteins. WT *C. albicans* more strongly suppressed secreted IL-6 and IL-12p40 in co-treated conditions than *Hgc1* KO (yeast locked) strain of *C. albicans*. This could suggest that phagolysosomal stress imparted by *C. albicans* hyphal transition may have an impact on selective cytokine suppression in BMDMs.

The growth of the hyphae puts stress on the phagosome, requiring merging of other host intracellular vesicles including lysosomes, delivering reactive oxygen species, hydrogen pumps lowering the pH and cathepsins as antimicrobial defences (77,78). Additionally, from within the phagosome, *C. albicans* has its own defence mechanisms including a pH neutralising processes and pore forming proteins which together enable *C. albicans* to escape the phagosome and then the host cell altogether (14,47).

Altogether, the data in this paper is consistent with the idea that *C. albicans* can act to evade macrophage activation and while also potentially acting to repress aspects of the macrophages inflammatory function in response to other inflammatory stimuli such as LPS or *P. aeruginosa* infection. This suppression requires contact between the *C. albicans* and the macrophages; however, its molecular mechanism is not clear. It does, however, have implications for how the host may respond to polymicrobial infections.

## Materials and methods

### Mice

Mice were used for the isolation of bone marrow derived macrophages (BMDMs) at either the University of Dundee or University Aberdeen. 8 to 12 week old male C57Bl6/J mice used in Dundee were obtained from Charles River Laboratories (UK) and experiments approved by the Ethical Review and Welfare Committee at the University of Dundee. The use of *Ptpn1*^fl/fl^ mice at University of Aberdeen was approved by the Ethical Review Committee at the University of Aberdeen and performed in compliance with the United Kingdom Animal (Scientific Procedures) Act 1986, under UK Home Office project licence number PPL 70/8073. *Ptpn1*^fl/fl^ mice were on a C57Bl6/J background, and as these mice did not express Cre recombinase, they were regarded as wild type for the purpose of this study. The generation of the *Ptpn1*^fl/fl^ mice has been described previously (79). All animals were maintained in accordance UK and EU regulations and were maintained under specific pathogen free conditions in individually ventilated cages. Animals had free access to food and water (R&M1 SDS, Special Diet Services) and kept under a 12/12-hour light / dark cycle at 21°C (Dundee) or kept at 22-24°C (Aberdeen) with 45-65% humidity. Mice were sacrificed using a rising concentration of CO_2_ with additional confirmation of death by cervical dislocation.

### Murine bone marrow derived macrophage cell culture

On day 0, bone marrow was flushed with 10 – 20 ml of PBS from the femurs and tibias of a mouse and filtered through a 100 μm strainer and centrifuged at 400 *g* for 5-minutes. The cells were then re-suspended in 10 mL of L929 conditioned media, DMEM supplemented media containing (DMEM (Gibco, 11960), 100 Units/mL Penicillin and 100 μg/mL Streptomycin (Gibco, 15140-122), 1 mM sodium pyruvate (Gibco, Cat. 11360), 1% 100x non-essential amino acids (Gibco, 11140), 50 μM 2-mercaptoethanol, 2mM GlutaMAX (Gibco, 35050) and 10mM HEPES (Lonza, BE17-737E), 10% heat-inactivated FBS (Labtech, 91242), with the addition of 20% L929 conditioned media. To generate L929 conditioned media, confluent L929 cells were maintained in DMEM supplemented media. After 1 week media was removed and retained and cells cultured for a further week in fresh media. This media was then removed and combined in equal volumes with the media from week 1 to give the final conditioned media. Bone marrow derived cells were cultured for 7 days on bacterial grade plates (Thermo Scientific, 101R20) at 37°C and 5% CO2. After 7 days, BMDM were detached by scraping in PBS with 1% EDTA, counted using a hemacytometer and replated at a density of between 0.2 and 0.5x106 cells/ml of BMDM media on tissue culture treated plates (Geiner), and incubated overnight before use.

### Microorganisms

*C. albicans* clinical strain SC5314 was donated by Dr Katharina Trunk (Newcastle University). The yeast-mutant *Hgc1* KO strain WYZ12.2 was donated by Professor Neil A. R. Gow (University of Exeter). *P. aeruginosa* strain PA01, was donated by Professor Megan Burgkessel (University of Dundee).

To culture *C. albicans*, a frozen stock of SC5314 *C. albicans* strain (20% glycerol, 80% YPD media containing *C. albicans* yeast) was used to streak a YPD agar plate (YPD medium + 2% agar) in sterile conditions and incubated overnight at 30°C. After distinct colony growth, a single colony was picked and cultured in 5 mL YPD media broth overnight in 30°C at 200 rpm. In the morning, the *C. albicans* was centrifuged at 400 *g* for 5 minutes, resuspended in 10 mL of PBS.

*C. albicans* was then counted on a hemacytometer. Volume was corrected with PBS to give 1x108 cells/mL, and then re-counted on the hemacytometer to confirm density. *C. albicans* was kept on ice until use for infection experiments on the same day as it was prepared. Heat-killed *C. albicans* were generated by treating *C. albicans* for 10-minutes at 100°C. Death was confirmed with streaking culture on YPD plate, overnight at 30°C. Sonicated *C. albicans*, had the additional step of sonication for 5 cycles of 30-second on, 30 seconds off at 50W in addition to the heat killing.

To culture *P. aeruginosa,* a frozen stock of *P. aeruginosa* strain PA01 (20% glycerol, 80% LB media containing *P. aeruginosa*) was streaked on LB plate and incubated at 37°C overnight. One day prior to *in vitro* experiment, a single colony was cultured in 5 mL of LB media broth overnight at 37°C and shaking at 200 rpm. The day of the experiment, 50 μL was removed from overnight culture and inoculated in 5 mL of fresh LB media and incubated for ∼2 hours at 37°C shaking at 200 rpm to return the bacteria to the log growth phase. The optical density of the culture was measured (OD600) using against LB media as a blank. PA01 Colony Forming Units (CFU) were then estimated as CFU/mL=OD600 x 7.39^^8^, based on previously generated standard curves for *P. aeruginosa* growth. Once the OD600 was measured, *P. aeruginosa* broth was centrifuged at 800*g* for 5 minutes and resuspended in PBS at a density that would give a MOI of 10:1 when diluted 1 in 1000 into macrophage cultures.

### Macrophage Treatments

BMDMs were infected with a multiplicity of infection (MOI) of 4:1 for *C. albicans* or 10:1 for *P. aeruginosa* unless otherwise stated. Plates were rocked manually back and forth to distribute the microorganisms, and then centrifuged at 150 g for 1-minute, to sediment the microorganisms. Plates were then placed in the incubator until ready for lysis and/or media collection for cytokine analysis. LPS from *Escherichia coli* O26:B6 (Sigma L2654) was used at 100 ng/mL. Where indicated in the figure legends, relevant PRRs were stimulated using either 1 µg/ml Pam3CSK4 (TLR1/2, Invivogen, tlrl-pms), 250ng/ml R848 (TLR7/8, Invivogen, tlrl-r848), 1 µg/ml CL097 (TLR7/8, Invivogen, tlrl-c97), 2μM ODN 1826 CpG Oligonucleotide (TLR9, Invivogen, tlrl-1826) or 200 µg/ml depleted Zymosan (Dectin1/TLR2, Invitrogen Tlrl-zyd).

### DIA-MS proteomic analysis

For proteomic experiments independent BMDM cultures from 4 C57Bl6/J mice were used. 1 million BMDMs per condition were lysed using 400 μL of 5% SDS (Sigma, 05030), 10mm TCEP (ThermoFisher Scientific, 77720), 50 mM TEAB (ThermoFisher Scientific, 90114) in HiPerSolv Water for HPLC (VWR, 83650.320). Lysis and proteomic sample preparation were as described (Baker *et al.*, 2022). Briefly, after lysis, lysates were incubated at 100°C for 5 minutes, and then sonicated before protein concentration was calculated using the EZQ protein quantification kit (Thermo Fisher Scientific, R33200). Tryptic peptides were generated by the S-Trap Method using S-Strap: Rapid Universal MS Sample Prep columns (Co2-mini, Protifi) and Trypsin Gold (Promega, V5280). Peptides were then vacuum dried and resuspended in 50 μL of 1% formic acid (Thermo Fisher Scientific, 695076). Peptide amounts were quantified via pierce Quantitative Fluorometric peptide Assay (Thermo Scientific 23290) prior to MS-analysis.

Samples were analysed on an Orbitrap Exploris 480 (ThermoFisher) coupled with a Dionex Ultimate 3000 RS (Thermo Scientific) in DIA mode, with 140min Grad-DIA. For each sample, 1.5 *μg* of peptide was analysed on the Exploris. Two blanks were run between each sample.

LC buffers prepared as follows: buffer A (0.1% formic acid in Milli-Q water (v/v)) and buffer B (80% acetonitrile and 0.1% formic acid in Milli-Q water (v/v)). 1.5 μg aliquots of each sample were loaded at 10 μL/minute onto a trap column (100 μm × 2 cm, PepMap nanoViper C18 column, 5 μm, 100 Å, Thermo Scientific) equilibrated in 0.1% trifluoroacetic acid (TFA). The trap column was washed for 5 min at the same flow rate with 0.1% TFA then switched in-line with a Thermo Scientific, resolving C18 column (75 μm × 50 cm, PepMap RSLC C18 column, 2 μm, 100 Å). The peptides were eluted from the column at a constant flow rate of 300 nL/min with a linear gradient from 3% buffer B to 6% buffer B in 5-minutes, then from 6% buffer B to 35% buffer B in 115 min, and finally to 80% buffer B within 7-minutes. The column was then washed with 80% buffer B for 6 min and re-equilibrated in 3% buffer B for 15-minutes. Two blanks were run between each sample to reduce carry-over. The column was kept at a constant temperature of 50°C at all times. The data was acquired using an easy spray source operated in positive mode with spray voltage at 1.9 kV, the capillary temperature at 250°C and the funnel RF at 60°C. The MS was operated in data-independent acquisition (DIA) mode. A scan cycle comprised a full MS scan (m/z range from 350–1650, with a maximum ion injection time of 20 MS, a resolution of 120,000 and automatic gain control (AGC) value of 5 × 10 6). MS survey scan was followed by MS/MS DIA scan events using the following parameters: default charge state of 3, resolution 30.000, maximum ion injection time 55 MS, AGC 3 × 106, stepped normalized collision energy 25.5, 27 and 30, fixed first mass 200 m/z. The inclusion list (DIA windows) and windows widths are published in Baker *et al*. 2022. The mass accuracy of individual peak spectra was checked prior the start of samples analysis.

### Spectronaut search and FASTA files

Raw data files for the 2 experiments were together in one search in Spectronaut 19 with the following FASTA files: *Mus musculus* SwissProt canonical with isoforms (November 2023), and *C. albicans* TrEMBL (May 2023).

All details of data file preparation can be found at (80). Optimised Spectronaut identification settings were used as reported in (81); briefly: precursor *q*-value cut-off 0.01, precursor PEP cut-off 0.01, protein FDR strategy accurate, protein *q*-value cut-off (experiment) 0.01, protein *q*-value cut-off (run) 0.01 and protein PEP cut-off 0.01 were used. The Proteomic Ruler was used to estimate protein concentration per cell (82,83) using Perseus (https://maxquant.net/perseus/) as well as the proteomic ruler plugin (http://www.coxdocs.org/doku.php?id=perseus:user:plugins:store).

All raw files, Spectronaut reports, FASTA files and experiment templates have been uploaded to PRIDE (84), under accession number PXD062689.

### Proteomic statistical tests

Prior to analysis of M*us musculus* proteins, *Candida albicans* proteins were removed from the dataset (Supplementary Table 7). The Bioconductor package, Limma (85) in R was used for statistical hypothesis testing for each proteomic comparison with linear models applied to each protein in a row producing *q*-value (also termed adjusted p-value ) and fold-changes. Differentially expressed proteins were defined using a thresholds of a *q* < 0.05 and a Log_2_ fold change of greater than 1 standard deviation away from the median. The robust list of differentially regulated proteins discussed in the text additionally were filtered, where proteins were only defined as up-regulated if present in ≥ 3 of the stimulated replicates, or if no proteins were present in the unstimulated yet present in 4 replicates in the treated conditions; and down-regulated if present in greater than ≥ 3 of the unstimulated replicates, or if present in 4 replicates of unstimulated, and not present in the untreated. Heatmaps were generated using Morpheus Heatmaps (https://software.broadinstitute.org/morpheus/).

### Immunoblotting

BMDMs are seeded in a 6-well tissue culture treated plate, 0.5 x10^6^ cells/well or equivalent density, and treated as described in the figure legends. Cells were washed once with PBS and lysed at room temperature with lysis buffer (50mM Tris-HCl, pH 7.5, 1% SDS (w/v), 1% (v/v) Triton-X-100 (ThermoFisher, 93443), 10% glycerol, 1mM EGTA, 1 mM EDTA, 1mM Sodium Orthovanadate, 50 mM Sodium Fluoride, 1 mM Sodium Pyrophosphate, 10 mM Sodium B-Glycerophosphate, 0.27 M Sucrose, 0.1% (v/v) 2-B-mercaptoethanol, complete mini EDTA-Free Protease Inhibitor tablet (Sigma, M6250)). Lysates were heated at 100°C for 5 minutes then frozen at -20°C. Before use, samples were thawed at room temperature and then passed 10 times through a 25 Gage BD MicrolanceTM 3 needle to sheer the DNA. Samples were run on 10% Tris-glycine gels using standard techniques. Proteins were then transferred onto nitrocellulose and membranes blocked in 5% milk in TBS-T (0.05M Tris-HCl pH7.6, 0.15M NaCl, 0.1% Tween-20). Primary antibodies were from Cell Signalling, listed in Supplementary Table 6. Detection was with an HRP-conjugated anti-rabbit secondary (Fisher, A16110) and Clarity Western ECL Substrate (BIO-RAD, 170-5061) and results visualised using a LI-COR Odyssey Fc Imager (LI-COR), with Image Studio Software. Full images of the membraneS are shown in Supplementary Figures 8 and 9.

### Cytokine Secretion Analysis

In order to measure cytokine secretion, media was collected from plates at the times indicated in the figure legend and stored at -20°C until analysis. Levels of IL-6, IL-10, IL-12p40, TNF, were determined using Bioplex Pro TM Mouse Cytokine Assays (Bio-Rad) according to the manufacture’s protocols. Data was acquired on Luminex-200 multiplexing instrument and analysed using xPONENT acquisition and analysis software (Luminex).

### Live/dead Imaging assay

BMDMs were plated as listed for a 6-well plate and treated as described in the relevant figure legend. After the treatment was complete, 50 nM Sytox green was added to media, and left in the dark for 5-minutes. Wells were then washed 5 times with PBS and then cells were fixed with 1:1 IC fixation buffer (Fisher, 00-8222-49 using 400 μL/well) and left at 4°C for 20-minutes. Cells were washed 3-times in PBS and then permeabilized using 400μl of 1x permeabilization buffer (Fisher, 00-8333-56) with 1.25 μg/mL DAPI for 10-minutes at room temperature. Images were then taken in green and blue field on a ZOE BIORAD microscope. Cell counts were done manually.

### Transwell assays

To determine the direct contact between the macrophage and *C. albicans* was important, assays were carried out in 6.5 mm Transwell® plates with inserts containing 0.4 µm pores. BMDMs were seeded in the lower part Transwell® plates overnight. The following day, *C. albicans* was added either straight onto BMDMs or into the centre of the Transwell insert. For co-treated conditions, 100 ng/mL of LPS was added directly onto BMDMs. BMDMs were left for 8-hours and then media was collected and Luminex cytokine assay was performed as described above. Cultures were kept overnight in 37°C and 5% CO_2_ incubator after stimulation to confirm no *C. albicans* growth was observed in wells (with BMDMs) where *C. albicans* was separated by the insert.

### Statistical analysis

For the analysis of individual proteins or Luminex data the statistical tests used are indicated in the legends. For analysis involving more than one group one way ANOVA or two way ANOVA were used. Analysis was carried out using Graph Pad Prism.

## Author Contributions

Conceptualization, JSCA, CPB, SL; methodology, JSCA, CPB, SL, JW, KS; formal analysis, CPB, SL; writing—original draft preparation and writing CPB; reviewing and editing, CPB, JSCA, JW, SL, HW. All authors have read and agreed to the published version of the manuscript.

## Data Availability Statement

Mass spectrometry data has been deposited in PRIDE (84), under accession number PXD062689. Full scans of immunoblots are available in the supplementary data. Other data is available on request from the authors.

## Acknowledgements

The authors would like to thank members of the Arthur and McSorley Lab for discussions pertaining to the figures and story of this paper; A. Brenes and A. Howden for proteomic analysis advice; Dundee University Biological Resource unit; A. Score, A. Atrih and the Fingerprint Proteomics Facility.

